# LbCas12-mediated multiplex gene editing and 2-fluoroadenine counter-selection in *Phytophthora palmivora*

**DOI:** 10.1101/2024.02.13.580060

**Authors:** Tim Verhoeven, Max HJ Pluis, Maaria Peippo, Gabriel Couillaud, Grardy CM van den Berg, Edouard Evangelisti

**Author notes:** Equal contribution (Tim Verhoeven, Max HJ Pluis).

## Abstract

CRISPR-Cas systems have moved forward genetic engineering in virtually any organism amenable to genetic modification. In particular, these systems have unlocked unprecedented possibilities to generate mutants in oomycetes, a group of filamentous microbes comprising over two hundred *Phytophthora* species, including the cacao killer *Phytophthora palmivora*. Here, we showcase multiplex gene editing in *P. palmivora* using LbCas12. We have developed a straightforward protocol to simultaneously knock out two genes encoding adenine phosphoribosyltransferase (APT), an essential enzyme of the purine salvage pathway. We show that *APT* knockouts (Δ*PpATP1/2*) are insensitive to 2-fluoroadenine (2-FA) and retain full virulence on *Nicotiana benthamiana*. We rely on zoospore electroporation using an all-in-one construct to facilitate the rapid editing of multiple genes. This work enhances the genetic toolbox for *Phytophthora* species and simplifies the exploration of gene function, laying the groundwork for future innovations aiming to tackle oomycete plant diseases.

## Introduction

Over the past decade, the prokaryotic Clustered Regularly Interspaced Short Palindromic Repeats (CRISPR)-Cas immune system has been repurposed for genome editing, moving forward genetic engineering (Wang and Doudna, 2023). Compared to protein-based techniques such as zinc finger nucleases (ZFNs) (Urnov et al., 2010) and Transcription Activator-Like Effector Nucleases (TALENs) (Sun and Zhao, 2013), CRISPR-Cas is more straightforward, adaptable, and cost-effective. The first widely adopted system, CRISPR-Cas9, originates from the bacterium *Streptococcus pyogenes*. It functions as a three-component ribonucleoprotein comprising the CRISPR-associated endonuclease Cas9 and two types of RNAs: a CRISPR RNA (crRNA) containing a 20-nucleotide (nt) spacer for sequence-specific targeting and a trans-activating CRISPR RNA (tracrRNA) (Jinek et al., 2012). Once bound to DNA, Cas9 cleaves DNA 3-4 nt upstream of the protospacer adjacent motif (PAM), a Cas-specific short nucleotide sequence (5’-NGG-3’ for SpCas9) essential for target identification (Jinek et al., 2012). The CRISPR-Cas9 system was refined by fusing crRNA and tracrRNA into a single guide RNA (sgRNA) using a short loop, forming a hairpin-like structure. sgRNAs can be conveniently expressed under the control of RNA polymerase (Pol) III promoters, which transcribe non-coding RNAs without adding a 5’ cap or a poly(A) tail (Dieci et al., 2007).

Cas nucleases with properties distinct from Cas9 have been identified. These include Cas9 nickases (Cas9n), which induce single-strand breaks to minimize off-target effects compared to standard Cas9 (Ran et al., 2013; Trevino and Zhang, 2014), and the RNA-targeting enzyme Cas13 (Shmakov et al., 2015). Another notable Cas enzyme, Cas12a, has dual nuclease activities, including an endoribonuclease activity within its wedge (WED) domain. This unique feature enables Cas12a to process its own pre-crRNA into a mature guide RNA (Paul and Montoya, 2020; Zetsche et al., 2015). Cas12a can thus process an RNA transcript encoding a crRNA array to achieve multiplexing, i.e., concurrently targeting multiple genomic sites (Campa et al., 2019; Liao et al., 2019; Zetsche et al., 2017). Multiplex gene editing based on CRISPR has been reported in various organisms, including plants (Wang et al., 2017), fungi (Swiat et al., 2017), and bacteria (Ao et al., 2018), and has great potential in gene therapy aiming to tackle diseases in humans (Bigini et al., 2023). While Cas12 has greater specificity and fewer off-target effects compared to Cas9, it also exhibits lower nuclease activity (Singh et al., 2018; Strohkendl et al., 2018) and recognizes a more specific PAM sequence (5’-TTTV-3’) than Cas9.

The need to effectively select CRISPR-Cas edited events arises from inherent challenges in distinguishing edited from unedited cells. One strategy involves knocking out the orotidine 5-phosphate decarboxylase (*UMPS*) gene, which encodes a key enzyme in the pyrimidine *de novo* synthesis pathway (Wiebking et al., 2020). *UMPS* knockouts are uracil-auxotrophs and exhibit resistance to 5-fluoroorotic acid (5-FOA), a precursor to toxic 5-fluorouracil (Koh and Wickens, 2014). 5-FOA thus enables the counter-selection of non-edited cells (Serif et al., 2018). Similarly, disrupting expression of the adenine phosphoribosyltransferase (*APT*) gene, which encodes an essential enzyme in the nucleotide salvage pathway (Gaillard et al., 1998), confers resistance to the adenine antimetabolite 2-fluoroadenine (2-FA) (Schaff, 1994). 2-FA treatment is used as an efficient counter-selection method in green algae and moss (lineage Viridiplantae) (Collonnier et al., 2017; Guzmán-Zapata et al., 2019; Ichihara et al., 2022; Kim et al., 2021) and in brown algae and diatoms (lineage Stramenopiles) (Badis et al., 2021; Serif et al., 2018). Consequently, *UMPS* and *APT* gene knockouts are valuable counter-selectable markers, facilitating the identification of successful CRISPR-Cas editing events.

Oomycetes are filamentous microbes that belong to the Stramenopiles clade alongside brown algae and diatoms (Beakes et al., 2012; Derelle et al., 2016). Many have a parasitic or pathogenic lifestyle, affecting diverse hosts, including algae, diatoms, insects, fish, mammals, and plants. Notable plant-pathogenic oomycetes include the late blight pathogen *Phytophthora infestans*, responsible for the Great Irish Famine and a persistent threat to the contemporary potato industry (Fry et al., 2015), the sudden oak death pathogen *Phytophthora ramorum*, which decimates oaks and larches (Grünwald et al., 2012), and *Phytophthora palmivora*, a pathogen of tropical crops like cacao, papaya, citrus, and oil palm (Torres et al., 2016). Successful CRISPR-Cas9 gene editing in oomycetes was first reported in the soybean root and stem rot pathogen *Phytophthora sojae* (Fang and Tyler, 2016). In the absence of a characterized Pol III promoter in oomycetes, a Pol II promoter was used to control the expression of the sgRNAs. sgRNA maturation was then mediated by hammerhead ribozyme sequences, enabling self-cleavage (Fang and Tyler, 2016). This protocol was proven to be applicable in other *Phytophthora* species (Fang and Tyler, 2016; Gumtow et al., 2018; Miao et al., 2018), except for *P. infestans*, a species that does not tolerate Cas9 (Ah-Fong et al., 2021; van den Hoogen and Govers, 2018). Conversely, Cas12a is not toxic for *P. infestans*. Successful gene-editing was achieved using *Lachnospiraceae bacterium*-derived Cas12a (LbCas12a) (Ah-Fong et al., 2021), and the gene-editing efficiency was further improved by introducing a D156R point mutation in LbCas12a to enhance enzymatic activity at 25°C (Mendoza et al., 2023).

This study establishes a straightforward pipeline to simultaneously edit multiple genes in *P. palmivora* using the CRISPR/Cas12 system and select edited lines. Using a time-efficient transformation protocol based on zoospore electroporation and the optimized LbCas12a^D156R^ variant, we showcase multiplex CRISPR editing by knocking out the two *P. palmivora* genes encoding APT. We further show that the *APT*-knockout lines (Δ*PpATP1/2*) are resistant to 2-FA treatment and retain virulence on *Nicotiana benthamiana* leaves. Our results pave the way for advanced genetic manipulation in a group of economically relevant plant pathogens.

## Results

### 2-fluoroadenine inhibits *P. palmivora* growth in a dose-dependent manner

To assess the potential of 2-FA for counter-selection in oomycetes, we examined zoospore germination and mycelium growth in *P. palmivora* cultured on V8 medium supplemented with increasing concentrations of 2-FA **(Figures 1, S1)**. Mycelium growth on plates containing 10 µM 2-FA was slightly reduced compared to the control, but sporangia were visible, suggesting sporangiogenesis was not altered at this concentration **(Figure 1)**. Conversely, higher concentrations of 2-FA resulted in complete growth inhibition **(Figure 1)**. Microscopic inspection of the plates revealed that cyst germination occurred at 50 µM 2-FA but was abolished at 100 µM, with inoculation spots containing ungerminated cysts only **(Figure 1)**. Growth inhibition in the presence of 2-FA was also observed in *P. infestans, Phytophthora capsici*, and *Phytophthora parasitica* **(Figure S1)**. Thus, 2-FA is a potent inhibitor of *Phytophthora* growth *in vitro*.

**Figure 1.**
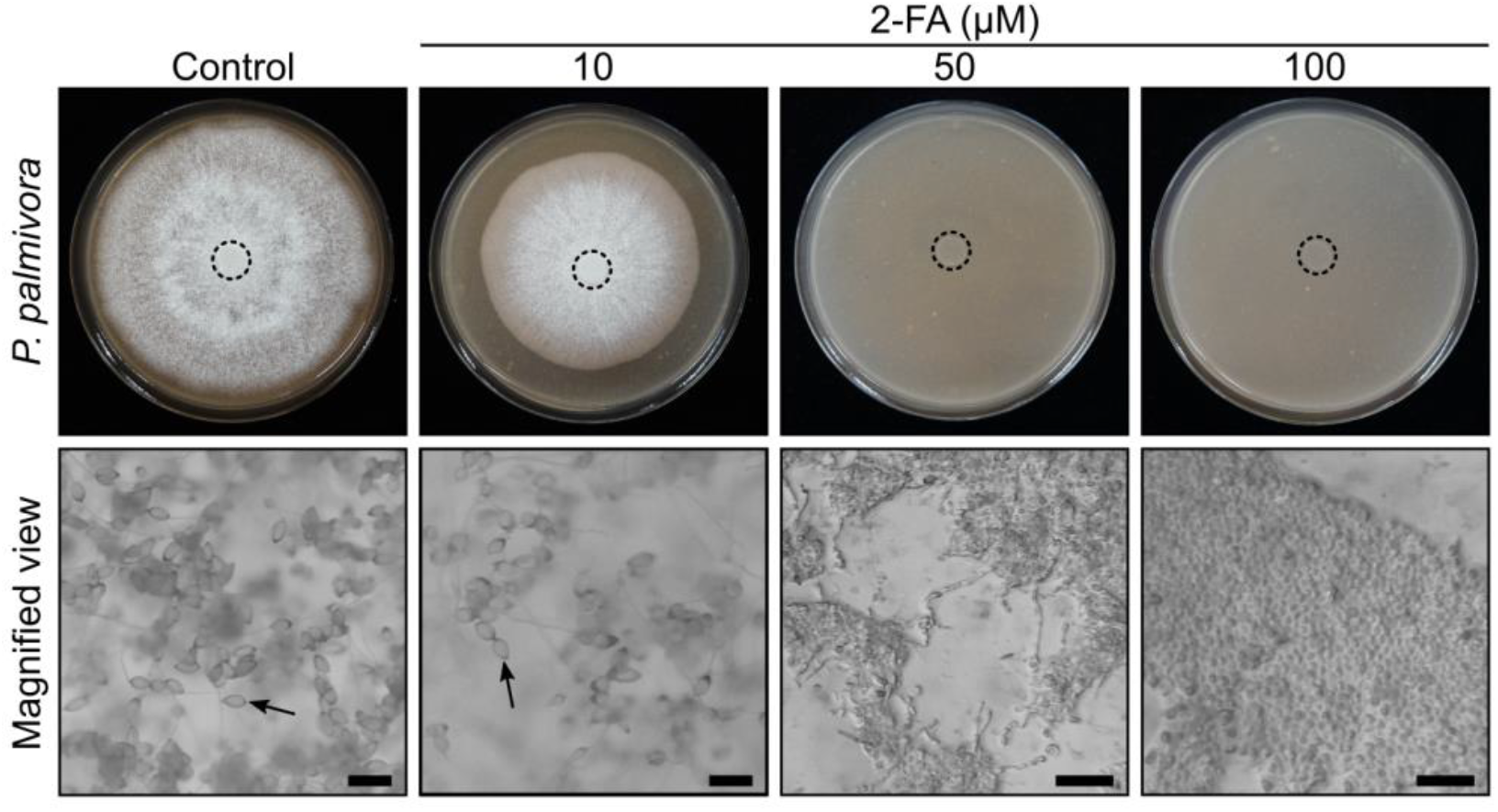
2-fluoroadenine inhibits *Phytophthora palmivora* growth in a dose-dependent manner. **Upper panels:** *P. palmivora* cultured on V8 medium without (control) or with 10, 50 or 100 µM 2-fluoroadenine (2-FA) for one week. Dotted circles indicate inoculation spots. **Lower panels:** magnified views of mycelial growth. Arrows indicate sporangia. Scale bars represent 100 µm.

### Two genes encode adenine phosphoribosyltransferase in *P. palmivora*

Next, we probed the *P. palmivora* genome (Ali et al., 2017) for genes encoding adenine phosphoribosyltransferase (APT). We found two gene models, *PHPALM_15494* (Genbank entry ID: POM68355.1), consisting of a single exon, and *PHPALM_36733* (Genbank entry ID: POM58597.1), containing three exons, and we named the genes *APT1* and *APT2*, respectively. Genes encoding proteins similar to APT1 and APT2 were found in other *Phytophthora* species **(Figure S2)**. Despite APT2 exhibiting 28% amino acid sequence identity with APT1 **(Figure 2a)**, both proteins showed a remarkable structural similarity with *Homo sapiens* APRT (PDB entry ID: 1zn8) when modelled with AlphaFold **(Figure 2b-d)**. Model confidence was high except in the N-terminus and flexible loops **(Figure 2b-d)**. The root-mean-square deviations (RMSD) between *P. palmivora* APT proteins and APRT were lower than 2 Å, indicating high structural overlap. In particular, the amino acids critical for substrate binding were conserved in the two proteins **(Figure 2)**. Hence, our findings show that *P. palmivora* has two genes encoding APT.

**Figure 2.**
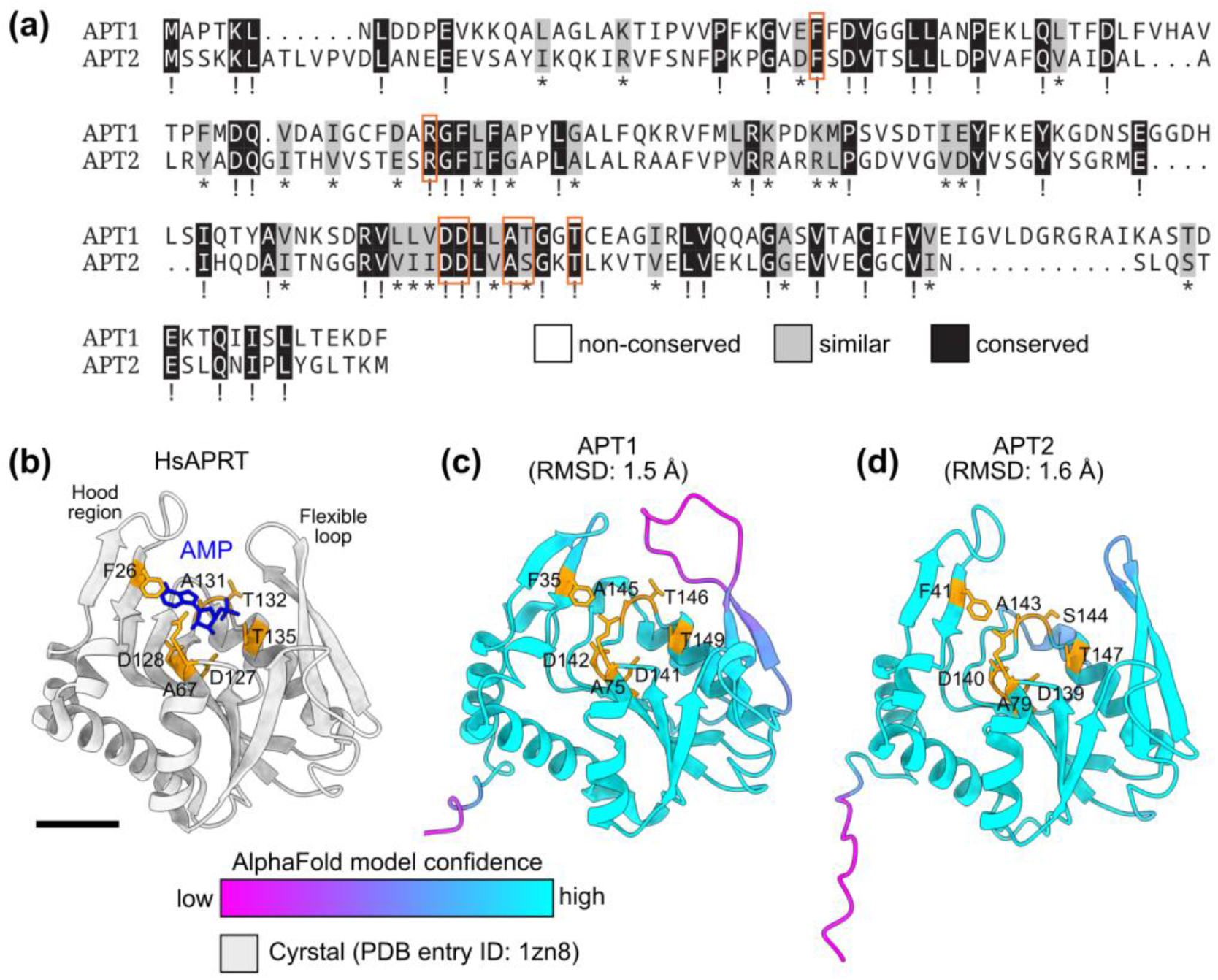
Two genes encode adenine phosphoribosyltransferase in *Phytophthora palmivora*. **(a)** Alignment of the deduced amino acid sequences of *P. palmivora APT1* (*PHPALM_15494*) and *APT2* (*PHPALM_36733*) with conserved and similar amino acids highlighted in black and grey, respectively. Orange frames indicate conserved residues located in the APT binding pocket. **(b)** Crystal structure of the *Homo sapiens* adenine phosphoribosyltransferase (HsAPRT, grey) complexed with adenosine monophosphate (AMP, blue) (PDB entry ID: 1zn8). The scale bar represents 1 nm. **(c-d)** Alphafold models of *P. palmivora* proteins APT1 **(c)** and APT2 **(d)**. Model confidence is color-coded. Conserved residues in the APT binding pocket are shown in orange. RMSD: root-mean-square deviation to HsAPRT.

### Disruption of *P. palmivora APT* genes confers resistance to 2-fluoroadenine

Next, we used the CRISPR/Cas12a system (Ah-Fong et al., 2021) to knock out the two *P. palmivora APT* genes at once **(Figure 3)**. First, we modified the pSTU-1 plasmid (Mendoza et al., 2023) by inserting a CRISPR array comprising three crRNAs **(Figure 3a)**. The first two crRNAs were designed to delete a 200 bp fragment in the *APT1* locus, while the third was designed to edit the *APT2* locus. Adjacent to the CRISPR array, we integrated a unique BsrG1 restriction site to facilitate further extension **(Figure 3a)**. We used the modified pSTU-1 plasmid for *P. palmivora* transformation via zoospore electroporation. Four rounds of transformations yielded 13 independent lines growing on geneticin (G418), indicating that the *nptII* gene located on pSTU-1 was successfully transferred to the recipient *P. palmivora* strain. Five lines (38%) showed sustained growth on V8 plates containing 2-FA **(Figures 3b, S3)**. With 2-FA in the medium, the colony sizes of lines 3, 8, and 10 were only slightly reduced **(Figure 3b)**, while lines 1 and 2 showed stunted growth even on V8-agar without 2-FA **(Figure S3)**, suggesting a possible position effect independent of gene editing.

**Figure 3.**
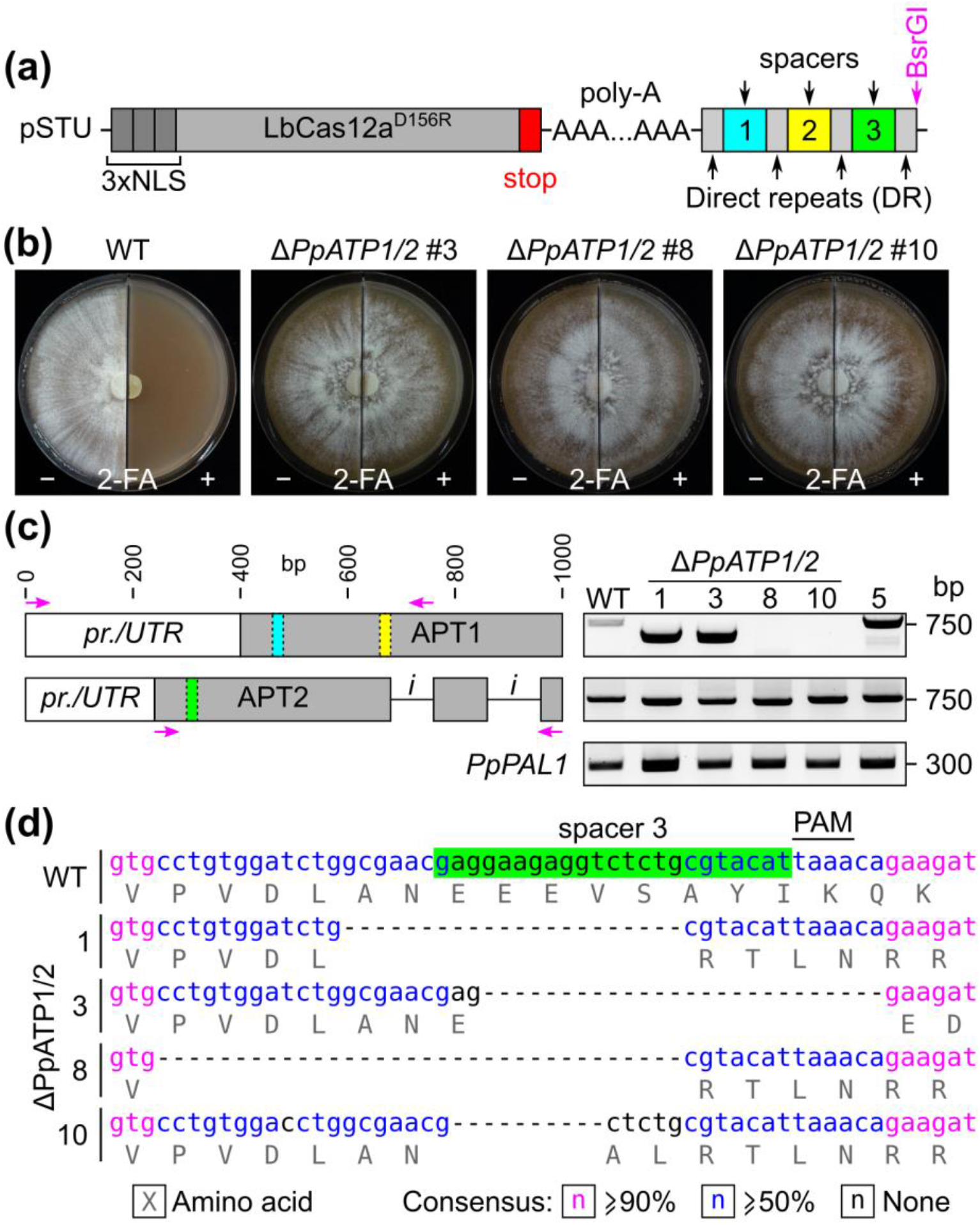
Disruption of *Phytophthora palmivora APT* genes confers resistance to 2-fluoroadenine. **(a)** Schematic representation of the modified pSTU-1 plasmid containing a crRNA array to knock out *P. palmivora APT* genes. The BsrGI restriction site (magenta) facilitates the further expansion of the array. **(b)** Growth of the recipient *P. palmivora* strain P16830 (WT) and *APT*-knockout (Δ*PpATP1/2*) lines 3, 8, and 10 on V8 medium without (left) or with 100 µM 2-FA (right) for one week. **(c)** Schematic representation of crRNA target sites in *APT1* and *APT2* (left) and the amplicons obtained by PCR (right) using the primers indicated by magenta arrows and DNA from wild-type (WT), Δ*PpATP1/2* lines 1, 3, 8, and 10, and G418-resistant line 5 as the template. Amplification of the *P. palmivora* elicitin gene *PAL1* (*PpPAL1*) is used as a control for the template DNA. *pr*.: promoter; *UTR*: 5’-UTR; *i*: intron. **(d)** *APT2* nucleotide sequences near spacer 3 (in green) in WT and Δ*PpATP1/2* lines 1, 3, 8, and 10. Uppercase letters (grey) indicate amino acids. Lowercase letters indicate nucleotides. Nucleotide consensus is color-coded. PAM: protospacer adjacent motif.

Next, we analysed 750-bp regions surrounding the crRNA binding sites by performing polymerase chain reactions (PCR) on genomic DNA from the three 2-FA-resistant lines 3, 8, and 10, and one of the two 2-FA-sensitive lines with stunted growth, line 1 **(Figure 3c)**. The wild-type (WT) strain and the 2-FA-sensitive line 5 were used as non-edited controls, while the *P. palmivora* elicitin gene *PAL1* (*PpPAL1*) was used as a control for PCR efficiency **(Figure 3c)**. Agarose gel electrophoresis revealed that *APT1* amplicons from the WT and line 5 matched the anticipated 750-bp size. Conversely, 500-bp fragments were obtained for lines 1 and 3, indicative of roughly 250-bp deletions due to double editing. No amplicon was detected for lines 8 and 10, suggesting larger deletions encompassing the reverse primer binding site. In contrast with *APT1*, amplicon size shifts were not detected in *APT2* **(Figure 3c)**. Amplicon sequencing confirmed the occurrence of short deletions and frameshifts that could not be resolved by agarose gel electrophoresis in the 2-FA-resistant lines (1, 3, 8, 10) **(Figure 3d, Supporting Dataset)**. Hence, the CRISPR/Cas12 system combined with zoospore electroporation enables the generation of mutants carrying multiple gene editing events in *P. palmivora*, leading to 2-FA resistance.

### *APT* knockout does not alter *P. palmivora* virulence on *N. benthamiana*

To assess the effect of *APT* knockout on *P. palmivora* virulence, we inoculated *N. benthamiana* leaves with zoospores harvested from wild-type *P. palmivora* and *APT*-knockout (Δ*PpATP1/2*) lines 3, 8, and 10 **(Figure 4)**. Lines 1 and 2 were excluded due to their aberrant growth phenotype **(Figure S3)**. We analyzed colonization by imaging chlorophyll autofluorescence four days after inoculation. Our observations revealed extensive leaf infection by the three Δ*PpATP1/2* lines **(Figure 4a-c)**, with no significant difference compared to the wild-type strain **(Figure 4d)**. Thus, *APT* knockout does not alter *P. palmivora* virulence in *N. benthamiana*, suggesting mutants lacking functional APT enzymes can be used for plant infection studies.

**Figure 4.**
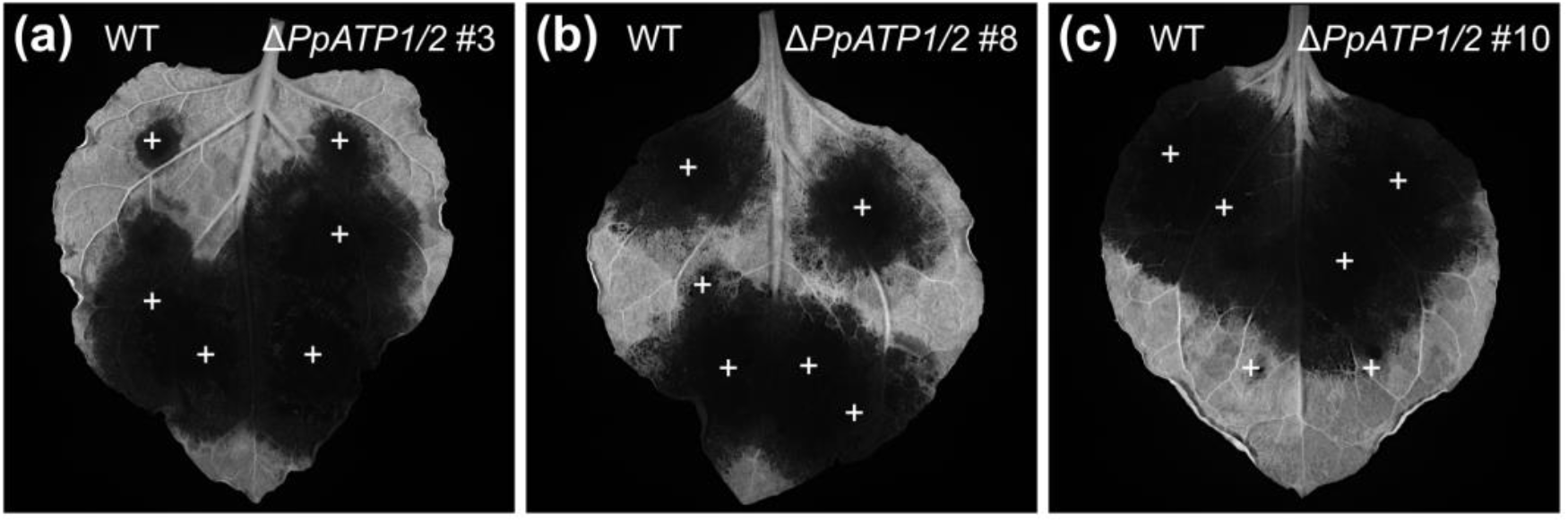
*APT* knockout does not alter *Phytophthora palmivora* virulence on *Nicotiana benthamiana*. **(a-c)** Representative images of *N. benthamiana* leaves infected with *P. palmivora* zoospores. Chlorophyll autofluorescence was observed four days after inoculation with zoospores from wild-type or *APT* knockout (Δ*PpATP1/2*) lines #3 **(a)**, #8 **(b)**, and #10 **(c)**. White crosses indicate inoculation sites.

## Discussion

We have identified two genes encoding adenine phosphoribosyltransferase (APT) in *P. palmivora* and established that *Phytophthora* growth is inhibited by 2-fluoroadenine (2-FA). *APT*-knockout lines maintained growth on plates supplemented with a concentration of 2-FA sufficient to inhibit zoospore germination (100 µM), which is notably higher than the concentrations used for counter-selection in *Ectocarpus* (20 µM) (Badis et al., 2021), *Physcomitrium patens* (10 µM) (Collonnier et al., 2017), and *Phaeodactylum tricornutum* (10 µM) (Serif et al., 2018). The lower sensitivity of *P. palmivora* to 2-FA may be due to differences in 2-FA uptake or purine metabolism in oomycetes. Furthermore, *P. palmivora APT* knockout lines did not require an exogenous adenine supply, a finding consistent with observations from *P. tricornutum* 2-FA-insensitive lines, which exhibited similar growth in the presence and absence of adenine (Serif et al., 2018). We also did not observe any change in the infection rate of *APT*-knockout lines in *N. benthamiana* leaf inoculation assays, i.e., in a context where pathogen access to nutrients is limited. Our findings align with the role of APT in the purine salvage pathway, a metabolic pathway independent of purine *de novo* synthesis (Ashihara et al., 2018; Meek et al., 2016). In contrast to *APT* knockout, knocking out *UMPS* leads to uracil auxotrophy (Sakaguchi et al., 2011), and this might impact virulence. Thus, *APT* knockout is more versatile and can speed up the selection of CRISPR mutants for plant infection studies.

Our study identified two genes encoding *APT* in *P. palmivora*. These genes encode proteins with divergent amino acid sequences but remarkable structural conservation, underscoring the benefit of 3D structure modelling over sequence-based analysis (David et al., 2022; Jumper et al., 2021). The occurrence of two *APT* genes in *P. palmivora* did not impede the practicability of this marker for counter-selection of non-edited cells. Although the number of transformants analysed in this study does not allow for a precise estimation of editing efficiency, our data suggest a slightly higher efficiency on two genes (38%) than previously reported on *P. infestans* using multiple crRNAs targeting a single locus (11-22%) (Mendoza et al., 2023). The difference in growth temperature between *P. infestans* and *P. palmivora*, the latter being maintained at a temperature where LbCas12^D156R^ activity is increased, may account for the higher efficiency. Besides, *P. palmivora* has a temperature maximum of 34°C (Puig et al., 2018), suggesting editing efficiency could be further increased, as higher editing efficiency was achieved with Cas12 enzymes at 28°C in plants (Malzahn et al., 2019) and in *P. infestans* (25%) (Mendoza et al., 2023). Overall, this showcases some advantages of *P. palmivora* as a model for genetic manipulation in oomycetes.

Combining 2-FA selection with zoospore electroporation allows for quick and efficient gene editing in *P. palmivora* and possibly other species amenable to zoospore electroporation, such as *P. capsici* (Huitema et al., 2011) and *P. infestans* (Dong et al., 2015; Latijnhouwers and Govers, 2003). Indeed, zoospores always contain a single nucleus (Walker and van West, 2007), lowering the risk of generating chimeric lines with heterogeneous nuclei populations. In contrast, oomycete protoplasts can be multinucleated (Schafrick and Horgen, 1978). In that case, heterogeneous nuclear subpopulations may occur and require the time-consuming isolation of single zoospore cultures (Ho and Ko, 1997). The occurrence of unedited nuclei in hyphae would result in APT being synthesised, resulting in growth inhibition on plates supplemented with 2-FA. Therefore, carefully selecting single spore derivatives is essential when combining 2-FA counter-selection with transformation methods other than zoospore electroporation.

We used G418 antibiotic resistance to select transformants and assess the frequency of *APT* gene editing. However, edited lines can be directly selected on 2-FA, removing the need for antibiotic selection and speeding up the process. For instance, a direct 2-FA selection was used to select *P. tricornutum* knockouts (Serif et al., 2018). However, this study relied on the biolistic delivery of CRISPR-Cas9 ribonucleoproteins, ensuring the direct availability of functional CRISPR complexes. However, the precise time frame for Cas12 expression and gene editing during plasmid-based transformation is unknown (Ah-Fong et al., 2021), and it remains to be tested if a direct 2-FA selection is feasible in our experimental setup.

## Conclusion

This study establishes an efficient multiplex CRISPR-Cas12a editing protocol for oomycetes, incorporating 2-FA-based counter-selection to accelerate the identification of editing events. Our approach will facilitate the study of multigene families by enabling scalable editing of multiple genes. The practicability of 2-FA selection paves the way for an alternative to antibiotic selection, offering more flexibility for subsequent complementation by super-transformation. Overall, the protocol we present here is a valuable addition to the genetic engineering toolkit for oomycetes and is expected to accelerate the unravelling of virulence and pathogenicity of *Phytophthora*.

## Experimental procedures

### Strain and growth conditions

*P. palmivora* Butler isolate LILI (accession number P16830) was initially isolated from African oil palms in Colombia (Torres et al., 2010). The strain is maintained on V8-agar in unsealed Petri dishes at 25°C under constant light. Transformed strains are maintained on V8-agar containing geneticin (G418) at 100 mg/L. For zoospore production, 1-week-old plates are incubated at 4°C for 30 min. Plates are then immersed with 10 ml sterile water and incubated at room temperature for 15 min.

### crRNA design

crRNAs were designed based on *P. palmivora* sbr112.9 genomic sequences (GenBank accession numbers: NCKW01008248.1 and NCKW01020197.1) (Ali et al., 2017). Three spacers **(Table S1)** were selected adjacent to the 3’ side of a 5’-TTTV-3’ PAM sequence. The first two spacers were designed to delete at least 200 bp of *APT1*, including the region encoding the binding pocket, while the third spacer targeted *APT2*. Sequences were checked for off-targets against the *P. palmivora* genome (Ensembl entry ID: ASM291172v1) using the CRISPOR software (http://crispor.tefor.net/) (Concordet and Haeussler, 2018). The CRISPR array extended with a BsrG1 restriction site at the 5’ end was synthesized in pUC19 (GenScript Biotech, Netherlands).

### pSTU-1 plasmid modification

The pSTU-1 plasmid carrying the optimized LbCas12a^D156R^ (Mendoza et al., 2023) was obtained from Howard Judelson. A fragment containing the CRISPR array plus the extra BsrG1 restriction site was cut from the synthesized pUC19 by digestion with DraI and BstZ17I enzymes and cloned into the BsaI-linearized pSTU-1 plasmid using the NEBuilder HiFi cloning kit (New England Biolabs, UK). The plasmid sequence was confirmed by sequencing (Eurofins Genomics, Germany).

### Generation of transgenic *P. palmivora*

*P. palmivora* transformation was carried out by electroporation of zoospores as previously described (Evangelisti et al., 2019) but with a higher zoospore concentration, approximately 108 zoospores/ml. Transformants were selected on 20% clarified V8-agar Petri plates containing 100 mg/L of G418 and transferred to fresh plates 5 days after electroporation.

### 2-FA sensitivity assays

For 2-FA sensitivity assays, 10 µL droplets of a *P. palmivora* zoospore suspension with 106 zoospores/mL were inoculated on V8-agar plates without and with 2-FA at final concentrations of 10, 50, or 100 µM. For growth assays on split Petri plates, circular mycelium plugs were cut with a cork borer from a 1-week-old mycelium culture and divided into two halves. The two halves were placed in the middle of the split Petri dish, one half on V8-agar without 2-FA and the other half on V8-agar containing 2-FA at a final concentration of 100 µM.

### Genotyping of 2-FA-resistant transformants

Primers outflanking crRNA binding sites were used to amplify *APT* coding sequences using Phire Plant Direct PCR Master Mix (Thermo Fisher, NL) according to the manufacturer’s instructions. Primer sequences are provided in **Table S2**. Amplicons were separated by agarose gel electrophoresis and imaged using a Gel Doc Go Imaging System (Bio-Rad, NL). Amplicons were validated by sequencing (Eurofins Genomics, Germany).

### Infection assays on *N. benthamiana*

For infection assays, leaves detached from four-week-old *N. benthamiana* plants, grown as described previously (Evangelisti et al., 2017), were inoculated with 10 µl droplets of a *P. palmivora* zoospore suspension with 105 zoospores/mL on the abaxial side and incubated at 23°C with a 16h photoperiod. Disease progression was monitored after three days using a ChemiDoc MP Imaging system (Bio-Rad, NL), using an excitation wavelength of 625-650 nm and a 695/55 nm bandpass emission filter.

## Acknowledgements

We are grateful to Howard Judelson (UC Riverside, USA) for providing the pSTU-1 plasmid. We thank Francine Govers (WUR Phytopathology, NL) for commenting on a draft version of the manuscript and Tijs Ketelaar (WUR Cell Biology, NL) for helpful discussions.

## Authors’ contributions

GvdB acquired data. TV designed experiments, acquired and analyzed data. MHJP, MP, and GC acquired and analyzed data. EE acquired funding, designed experiments, acquired and analyzed data, and wrote the manuscript with inputs from TV and MHJP. All authors have read and approved the final version of the manuscript.

