## Supporting Information for "LbCas12-mediated multiplex gene editing and 2-fluoroadenine counter-selection in *Phytophthora palmivora*"

**Contents**

Figure S1. Effect of 2-fluoroadenine on axenic growth of *Phytophthora spp*. 2

Figure S2. Phylogenetic tree of adenine phosphoribosyltransferases in oomycetes 3

Figure S3. Growth habit of the transgenic *Phytophthora palmivora* lines used in this study 4

Table S1. Spacers used in this study 5

Table S2. Primers used in this study 5

**Note:** Chromatograms (Supplementary Dataset) are provided as a separate ZIP file

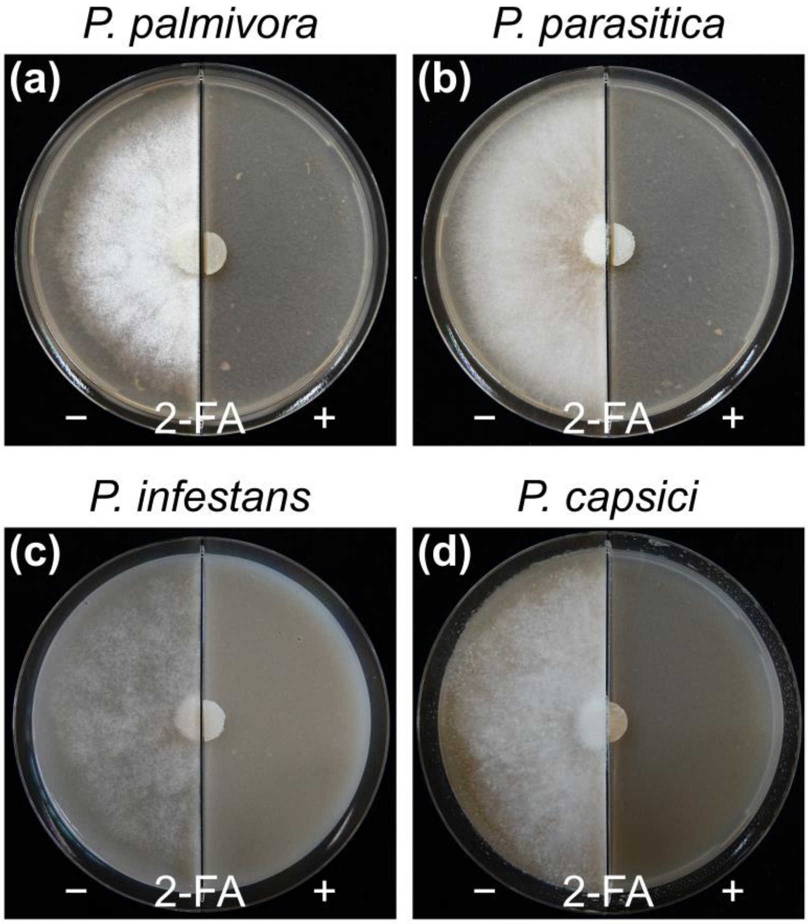


**Figure S1. Effect of 2-fluoroadenine on axenic growth of *Phytophthora spp*. (a-d)** Representative pictures of *Phytophthora palmivora* **(a)**, *Phytophthora parasitica* **(b)**, *Phytophthora infestans* **(c)** and *Phytophthora capsici* **(d)** grown on V8 medium, and *Phytophthora infestans* **(d)** grown on RSA medium, without (left) or with 100 µM (right) 2 fluoroadenine (2-FA).

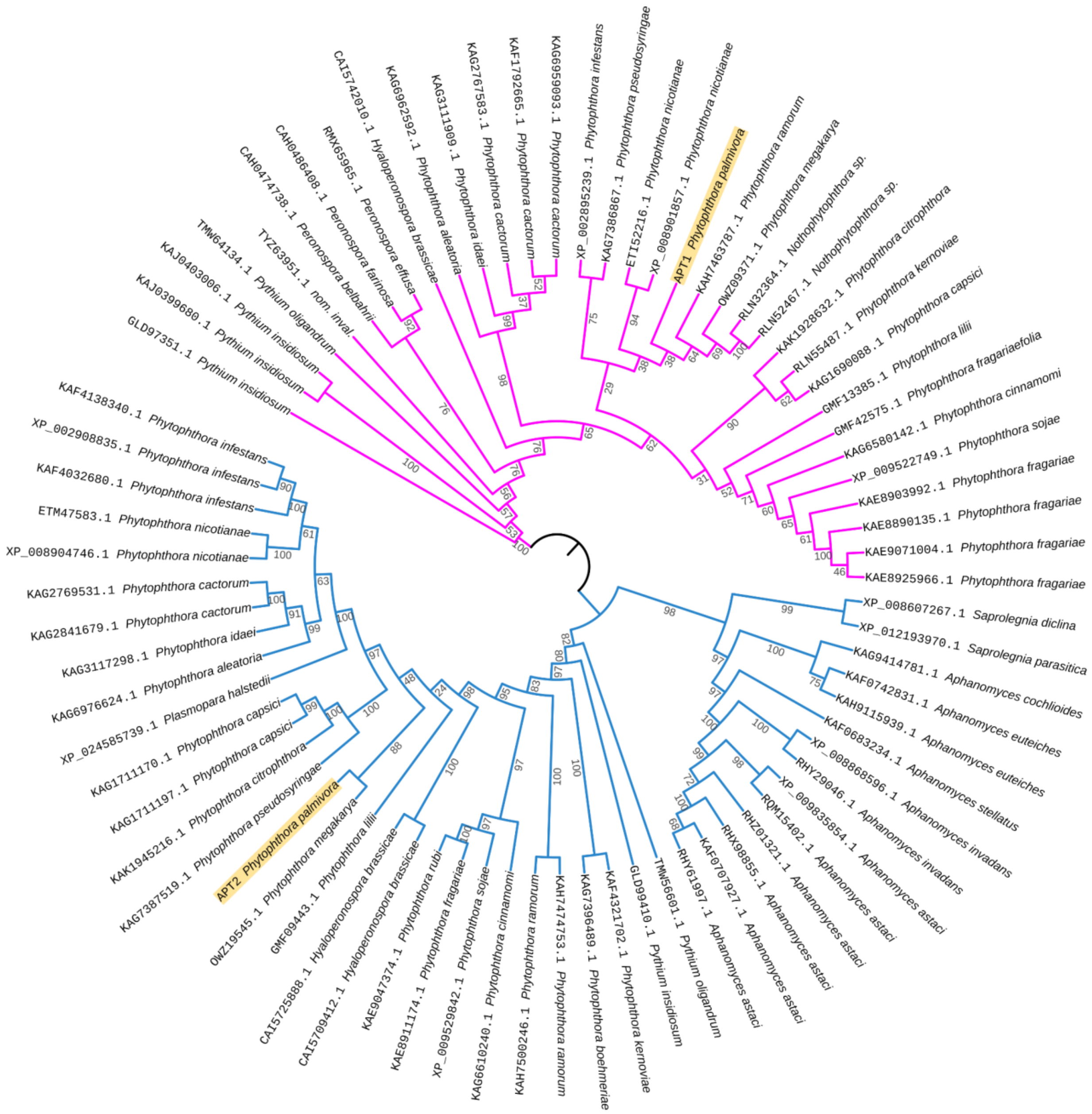


**Figure S2. Phylogenetic tree showing the evolutionary relationships of adenine phosphoribosyltransferases (APT) in oomycetes.** The Ultrafast Bootstrap method was used to build the phylogenetic tree using the IQ-TREE program (Hoang et al., 2018). *Phytophthora palmivora* APT1 and APT2 are highlighted in yellow. APT1 and APT2 subclades are displayed in magenta and blue, respectively.

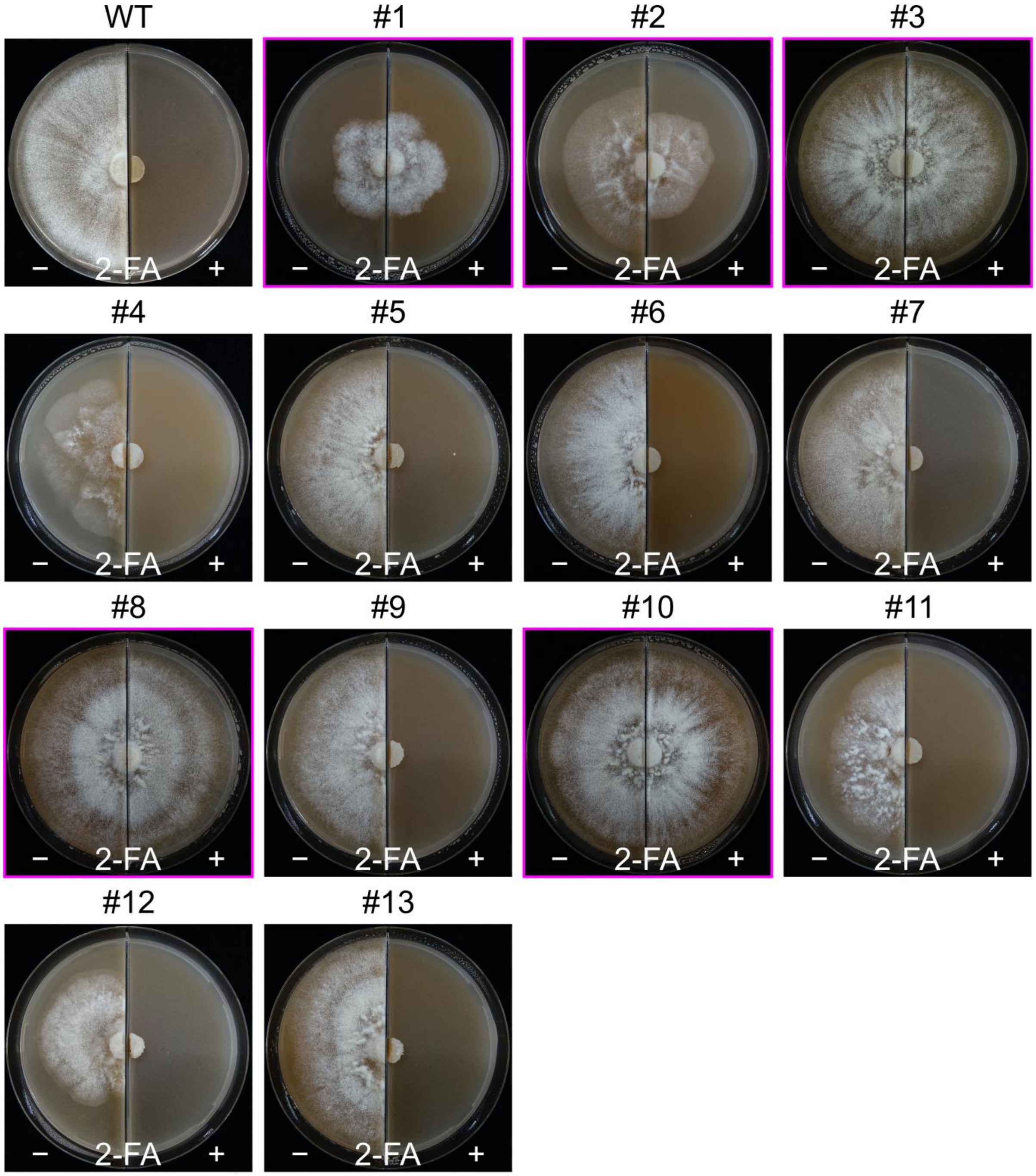


**Figure S3. Growth habit of the transgenic *Phytophthora palmivora* lines used in this study**, grown on V8 medium without (left) or with (right) 100 µM 2-fluoroadenine (2-FA). Magenta frames indicate 2-FA-insensitive lines.

**Table S1.** Spacers used in this study.

| Spacer name | Sequence (5’ → 3’) |
|---|---|
| Spacer 1 | AAAGGCACCACGGGGATGGTCTT |
| Spacer 2 | TCGGGTTTGCGCAGCATGAAGAC |
| Spacer 3 | ATGTACGCAGAGACCTCTTCCTC |

**Table S2.** Primers used in this study.

| Primer name | Sequence (5’ → 3’) |
|---|---|
| APT1_F | GGTGGTGGGTGGCTATGGATA |
| APT1_R | CCTCACTGTTGTCGCCCTTG |
| APT2_F | TGAGCAGCAAGAAGCTCGCAA |
| APT2_R | GCGGGATGTTCTGCAGAGATTCC |
| PpPAL1_F | ACCACGTGCACCACCACCCA |
| PpPAL1_R | TTATAGCGATGCACACGTAG |
